# Characterizing Solute Transport Across Cell Layers: Artifact Correction and Parameter Extraction from a Simplified Three-Compartment Model

**DOI:** 10.1101/2025.08.19.671057

**Authors:** Júlia Tárnoki-Zách, Imre Boldizsár, Gábor M Kovács, Balázs Döme, Szilvia Bősze, András Czirók

## Abstract

Quantifying solute transport across epithelial cell layers grown on transwell inserts is a common approach in early-stage drug development to estimate pharmacokinetic properties such as absorption and bioavailability. To increase throughput and reduce variability, these assays are increasingly automated, including the use of robotic or microfluidic systems for time-resolved sampling. However, both automated and manual sampling can introduce systematic artifacts, such as residual volume retention and surface adsorption, that distort concentration time series and affect downstream analysis. To fully realize the potential precision of automated measurements, we propose a mathematical correction to account for sampling artifacts; then to fit the corrected data to a three-compartment model that captures membrane diffusion, cellular sequestration, and metabolic loss. The method is demonstrated on datasets from transwell epithelial barrier transport assays. We suggest that the considered three-compartment model yields mechanistically more meaningful parameters than the conventional apparent permeability (Papp) measure. The proposed approach thus enables more accurate characterization of analyte interactions with the barrier cell layer, supporting better-informed assessments of compound behavior in in vitro transport systems.

## 1. Introduction

Permeability–the ability of molecules to cross biological barriers, is an important factor in drug development, influencing a compound’s absorption, distribution, metabolism, and excretion [1, 2, 3]. Promising drug candidates may fail due to inadequate permeability, which limits their ability to reach therapeutic concentrations at the target site [4, 5]. To assess this property, in vitro models have become indispensable tools for early-stage preclinical screening. Among these, cell-based barrier models, such as those employing Caco-2 [6] or MDCK [7, 8] cells cultured on transwell inserts [9], are particularly valuable [10, 11]. These models replicate several structural and functional features of biological barriers by forming confluent epithelial monolayers on porous membranes. Unlike artificial membranes, which model only passive diffusion, cell-based systems can capture a broad spectrum of biological processes, including active transport and facilitated diffusion, as well as intracellular metabolism and reversible sequestration. As these processes can influence drug disposition and bioavailability, cell-based barrier models can mimic physiological barrier functions more closely and often show good correlation with in vivo absorption data [6, 12, 13].

However, the advantages of cell-based barrier models come with increased complexity. The preparation of confluent, differentiated monolayers requires delicate handling in sterile facilities and extended culture times. Thus, epithelial barrier models are targets of automation efforts [14, 15, 16]. A typical assay involves time-resolved sampling of the basolateral (receiver) compartment following apical application of the analyte. While conceptually straightforward, this process is susceptible to experimental artifacts that, if uncorrected, can significantly bias estimates of kinetic parameters. In particular, the act of sampling - removing fluid from the receiver compartment and replacing it with fresh medium - can introduce distortions. Residual droplets retained in tubing or sampling hardware, as well as reversible adsorption of analytes to container surfaces, can blur temporal profiles, attenuate signals, or generate spurious peaks in the measured sample sequence. As we demonstrate, these artifacts disproportionately affect early time points, which are critical for resolving rapid transport processes such as membrane diffusion and cellular uptake.

In this work, we present an integrated experimental and modeling approach that addresses these challenges. First, we introduce a linear recursive model of the sampling process, which accounts for the two primary distortion mechanisms: residual volume retention and surface adsorption. This model is calibrated using cell-free experiments to estimate correction parameters under well-defined conditions. Next, we fit the corrected sampling data to a three-compartment model of passive transport across a cell-covered membrane. This model incorporates not only membrane permeability but also reversible analyte sequestration within the cell layer and irreversible intra-cellular loss (e.g., metabolic degradation). Together, these features allow the model to capture key aspects of analyte interaction with cellular barriers and to yield improved estimates of effective permeability. Finally, we determine under which conditions the model parameters can be reliably inferred from experimental data. This analysis reveals regions of parameter space where distinct parameter combinations produce similar outputs, highlighting the importance of careful experimental design and model validation. The presented approach enables more accurate estimation of permeability and associated kinetic parameters, helping to distinguish true biological transport phenomena from technical artifacts and thus making in vitro models more predictive and mechanistically informative.

## 2. Methods

### 2.1. Sampling apparatus

The analyzed measurement data were obtained using the Millitransflow an automated sampling system (BioPhys-Concepts, Budapest, Hungary; https://biophys-concepts.com), as described in [16]. The system employed four independent fluid lines: two for delivering fresh medium and analyte-containing medium, and two for removing fluid from the culture well and the apical compartment of the transwell insert. In the tissue culture incubator the multiwell plate was placed on a rotary shaker providing intermittent rotary motion at a radius of 10 mm and top rotational speed of 120 rpm.

### 2.2. Numerical methods

Data analysis has been performed using the scipy python module version 1.6.0. The analysis scripts are available at github. Ordinary differential equations were integrated using the solve_ivp function from the scipy.integrate package. Parameter optimization used the minimize function from the scipy.optimize package, using either the quasi-Newton method of Broyden, Fletcher, Goldfarb and Shanno (BFGS) [17] or the Nelder-Mead methods [18].

## 3. Results

### 3.1. Modeling and Correcting Sampling Artifacts

#### 3.1.1. Sources of sampling distortion

In automated sampling systems, the accuracy of the measured analyte amount depends on the quality of the sampling process. Unfortunately, various physical and chemical aspects of fluid handling can introduce systematic distortions that, if uncorrected, may compromise data interpretation. Here, we focus on two principal sources of such artifacts: (i) retention of residual liquid in the sampling path, and (ii) adsorption of analytes to surfaces in contact with the sampled fluid.

The sampling system we consider here typically includes a tissue culture chamber with a transwell insert, tubing segments, and sample vials. During each sampling cycle, a defined volume of fluid is aspirated from the transport compartment, transferred through the tubing, and deposited into a collection vessel. However, surface wetting often results in incomplete fluid displacement, leaving a residual volume within the system after the sample is delivered. In the absence of an intermediate wash step, this residual fluid is carried over into subsequent samples. Consequently, each collected sample contains carryover from previous cycles, introducing a temporal convolution of the true analyte sample profile. This distortion can appear as delays, attenuated slopes, and smoothing in the measured samples.

In addition to volume-related effects, small molecules − particularly hydrophobic or amphipathic compounds − may adsorb to plastic or glass surfaces along the sampling pathway. Such adsorption is typically reversible, with desorption occurring through prolonged contact with the fluid, potentionally over successive sampling cycles. This process also produces apparent delays, flattened slopes, and smoothing in the measured analyte profile, similar in effect to residual droplet carryover, but with a distinct underlying time dependence that reflects the dynamics of surface binding and release.

#### 3.1.2. Mathematical model of sampling distortion

To account for the above-discussed effects introduced during automated sampling, we developed a linear recursive model that captures the main mechanisms of sampling distortion: adsorption to surfaces and retention of residual droplets. This model builds on concepts introduced in an earlier work [16].

Let *S* = {*S*_1_, *S*_2_, …, *S*_*N*_} denote the measured analyte amounts in a sequence of *N* samples. Let *x*_*i*_ and *y*_*i*_ denote, respectively, the amount of analyte retained in residual droplets and adsorbed to internal surfaces immediately after the *i*th sampling cycle. Let *I*_*i*_ be the actual amount of analyte that entered the sampling (e.g., basolateral) compartment during the time interval between samples *i* − 1 and *i*. Due to sampling distortion, *S*_*i*_ ≠ *I*_*i*_, and the aim of the mathematical model is to relate these quantities.

Our sampling model is based on a balance calculation performed at each sampling cycle. Ve consider the total amount of analyte *T*_*i*_ that becomes available in the fluid phase of the sampling system during cycle *i*. This total reflects the incoming analyte from the transport process, *I*_*i*_, combined with carryover from the previous cycle: analyte retained in residual droplets (*x*_*i*−1_) and analyte desorbed from internal surfaces, modeled as *k*_*off*_ · *y*_*i*−1_. Thus, the total available analyte in sampling cycle *i* is

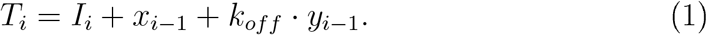

This total *T*_*i*_ is then partitioned into three fractions: (i) a portion reaches the sample vial and is recorded as the measured amount *S*_*i*_, (ii) a portion is adsorbed to internal surfaces and contributes to the updated surface-bound pool *y*_*i*_, and (iii) the remainder is retained in residual droplets, *x*_*i*_. The measured sample amount *S*_*i*_ is given by

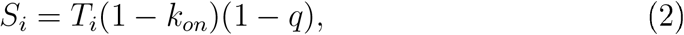

where *k*_*on*_ is the fraction of analyte lost to adsorption, and *q* is the volume fraction retained as residual liquid. The remaining analyte is distributed between adsorption and droplet retention according to

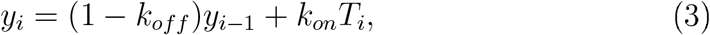

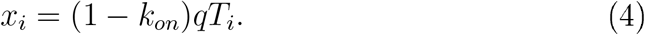

The linear recursive system of Eqs. (1) − (4) updates the internal states *x*_*i*_ and *y*_*i*_ at each sampling step and predicts the full sample sequence *S* = {*S*_1_, *S*_2_, …, *S*_*N*_} from a given transport inflow sequence *I* = {*I*_1_, *I*_2_, …, *I*_*N*_}.

To solve the inverse (deconvolution) problem, which involves recovering the transport inflow sequence *I* from a known measured sample sequence *S*, we employ a fitting process. Given the parameters *θ*_*sampling*_ = {*k*_*on*_, *k*_*off*_, *q*}, we generate candidate inflow sequences 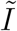 and calculate the corresponding predicted sample sequences 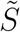 as defined by the sampling model. The best estimate of the inflow sequence, *I*, is identified as the candidate 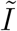 that minimizes the deviation

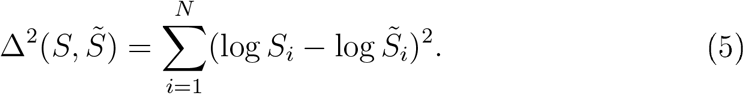

The comparison is performed on a logarithmic scale, as the analyte is often diluted by several orders of magnitude during the transport experiment.

#### 3.1.3. Two-compartment model for a cell-free membrane

The second model component describes the passive transport of the analyte across an inert, cell-free membrane separating two fluid compartments. The state variables are the concentrations in the apical and basal compartments, denoted *c*_*A*_(*t*) and *c*_*B*_(*t*), respectively. Their time evolution is governed by the conservation of mass:

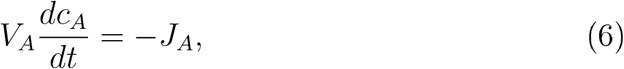

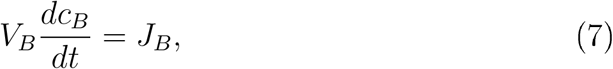

where *V*_*A*_ and *V*_*B*_ are the fluid volumes of the apical and basal compartments. For simple passive diffusion through a homogeneous, inert membrane, the flux across the membrane is given by Fick’s law:

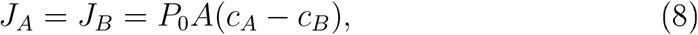

where *P*_0_ and *A* denote the permeability and surface area of the membrane, respectively.

To simulate a sampling measurement, the system of ordinary differential equations (6)–(8) is numerically integrated between discrete sampling time points. The simulation is initialized with *c*_*A*_(0) = *c*_0_ and *c*_*B*_(0) = 0. t the sampling time point of cycle *i*, the analyte content in the basal compartment is recorded as an element of the two-compartment inflow sequence, *Î*_*i*_, and *c*_*B*_ is reset to zero to mimic fluid removal by sampling. This procedure generates the model-predicted inflow sequence *Î* = {*Î*_1_, *Î*_2_, …, *Î*_*N*_}, where each element *Î*_*i*_ represents the total amount of analyte that diffused through an idealized membrane (assumed thin, homogeneous, and non-binding) between two successive sampling events.

To estimate the model parameters *θ*_*membrane*_ = {*P*_0_, *c*_0_} that best explain a given inflow sequence *I*, simulations are performed with various candidate parameter sets *θ*_*membrane*_. The best-fit parameter set is selected by minimizing the discrepancy between the given and model-predicted inflow sequences, Δ^2^(*I, Î*), using the same error metric defined in Eq. (5).

#### 3.1.4. Parameter estimation by nested fitting

To determine the sampling parameters *θ*_*sampling*_ = {*k*_*on*_, *k*_*off*_, *q*} that best characterize the distortion introduced during automated sampling, we implemented a nested fitting procedure (Fig. 2) that utilizes two types of experimental data: (i) transport experiments across cell-free membranes, and (ii) calibration experiments conducted in the absence of a transwell insert. These two experimental conditions impose complementary constraints on the estimation of sampling artifacts.

**Figure 1.**
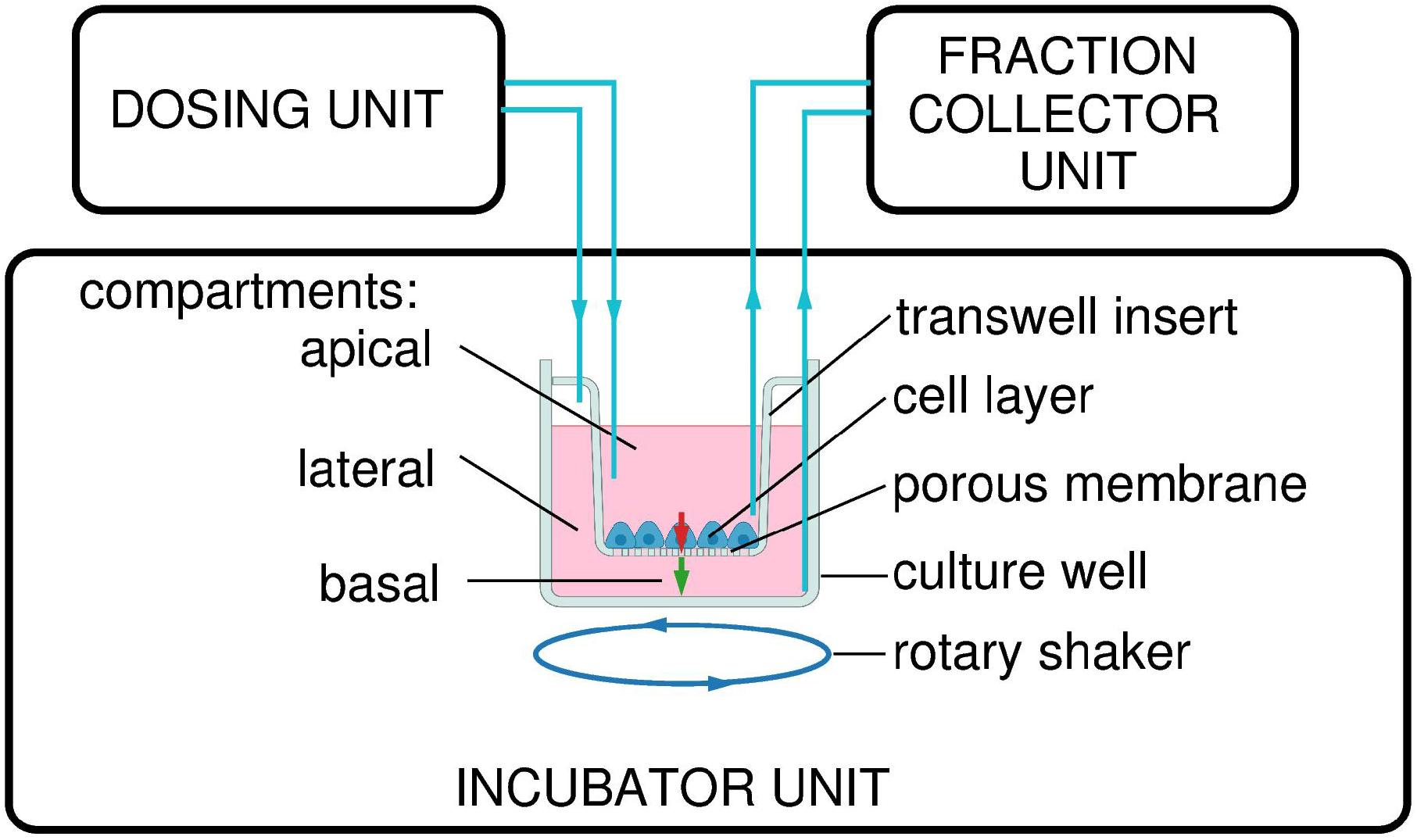
Diagram of the automated sampling system and incubator unit. Key components include the culture well with transwell insert (porous membrane and cell layer)’ apical’ basal’ and lateral compartments’ and a rotary shaker. Analyte fluxes *J*_*A*_ (apical efflux) and *J*_*B*_ (basal influx) are indicated by red and green arrows’ respectively. The dosing unit and cooled fraction collector are connected via four independent fluid lines for medium delivery and sample removal.

**Figure 2.**
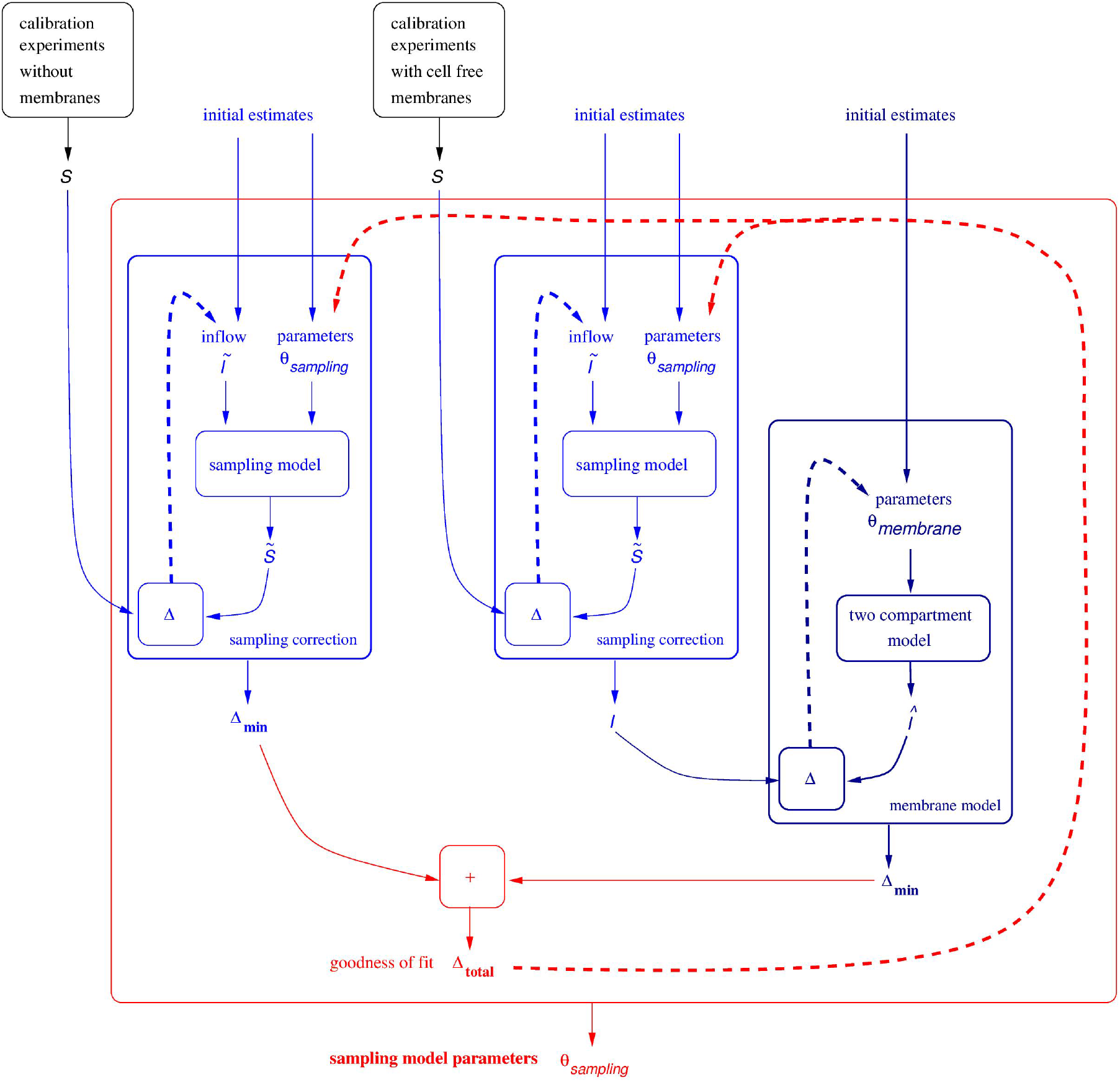
Nested fitting procedure for estimating sampling model parameters from calibration experiments, performed either with or without a cell-free transwell membrane. Boxes and arrows indicate algorithmic blocks with inputs and outputs. Dashed lines indicate optimization. For each candidate parameter set *θ*_*sampling*_, the sampling model (blue)is used to correct measured sample sequences *S* (black), yielding inflow sequences *I* and an associated goodness-of-fit estimate Δ_min_. For calibration experiments using a transwell membrane, the corrected inflow sequences *I* are used to fit the two-compartment membrane diffusion model (dark blue), which produces predicted inflow sequences *Î*. The discrepancy between *I* and *Î* contributes to the overall goodness of fit for the compound fitting procedure. The total objective function, defined as the sum of Δ_min_ across experiments, is minimized by tuning *θ*_*sampling*_, the parameters of the sampling distortion model. For readability, upper indices used in the main text for dataset indexing are omitted in the figure.

Let 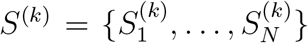 denote the measured sample sequences from independent experiments indexed by *k*. These datasets include experiments conducted both with and without a transwell membrane. For a given candidate parameter set *θ*_*sampling*_, we first apply the sampling distortion model to correct each measured sequence *S*^(*k*)^, yielding the corresponding inflow sequence *I*^(*k*)^ that estimates the true analyte delivery to the sampling compartment. For membrane transport experiments, each inflow sequence *I*^(*k*)^ is then fitted using the two-compartment passive diffusion model, resulting in best-fit membrane parameters 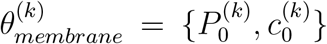 and a predicted inflow sequence *Î*^(*k*)^ derived from the idealized membrane transport assumptions. The discrepancy between the sampling-corrected inflow *I*^(*k*)^ and the two-compartment model prediction *Î*^(*k*)^ is quantified by the logarithmic de-viation Δ^2^(*I*^(*k*)^, *Î*^(*k*)^). For calibration experiments without a membrane, no transport model is required. In these cases, we compute the logarithmic deviation between the measured sample sequence *S*^(*k*)^ and the prediction 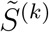 produced by the sampling distortion model, assuming a known inflow sequence *I*^(*k*)^ in which only the first element is nonzero.

The total fitting error across all datasets is the sum of transport model fitting errors and calibration model reconstruction errors:

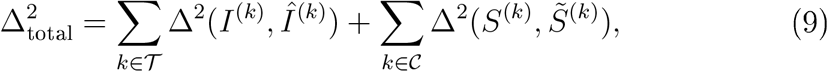

where 𝒯 and 𝒞 denote the sets of membrane transport and insert-free calibration experiments, respectively. As *I*^(*k*)^, *Î*^(*k*)^ and 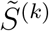 all depend of *θ*_*sampling*_, the condition (9) allows the determination of the optimal sampling parameter set as the one that minimizes 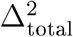.

#### 3.1.5. Sampling correction examples

To demonstrate the applicability of our sampling correction and diffusion modeling pipeline, we applied the method to two datasets. The first involved chloroquine (Fig. 3), a small-molecule compound known for its adsorption to various surfaces [19]. The second used fluorescein isothiocyanate−dextran (FITC-dextran, Fig. 4), a hydrophilic macromolecule that exhibits minimal binding to plastic surfaces. Both datasets included samples collected from the basolateral compartment of cell-free membranes, as well as from control experiments performed without a transwell insert. These datasets are used here solely to demonstrate the capabilities of the modeling pipeline. Their specific measurement methods and analytical protocols are described elsewhere [20].

**Figure 3.**
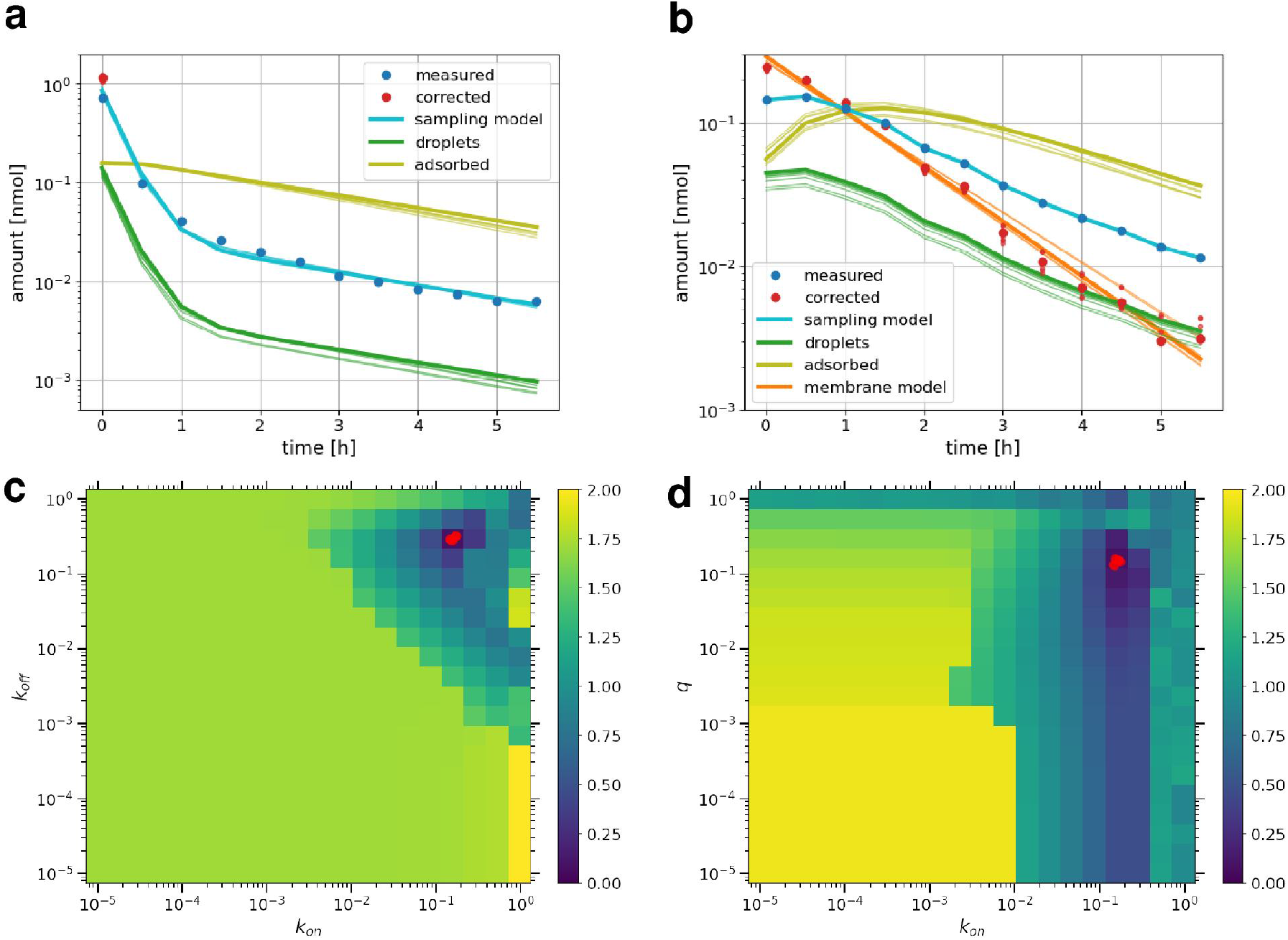
Sampling correction for chloroquine. (a) Measured sample sequence (blue), corrected inflow sequence (orange), and fitted sampling model (green) for membrane-free calibration experiments. Also shown are the predicted amounts retained in residual droplets (red) and adsorbed to surfaces (magenta), with jackknife uncertainty indicated by shaded traces. (b) Same as (a), but for diffusion across a cell-free membrane. The brown line shows the best-fit two-compartment model of passive membrane transport. (c) Heatmap of total fitting error Δ_total_ across the (*k*_on_, *k*_off_) parameter plane. The red dot indicates the best-fit parameter set smaller red points represent jackknife estimates. (d) Same as (c), shown across the (*k*_on_, *q*) plane.

**Figure 4.**
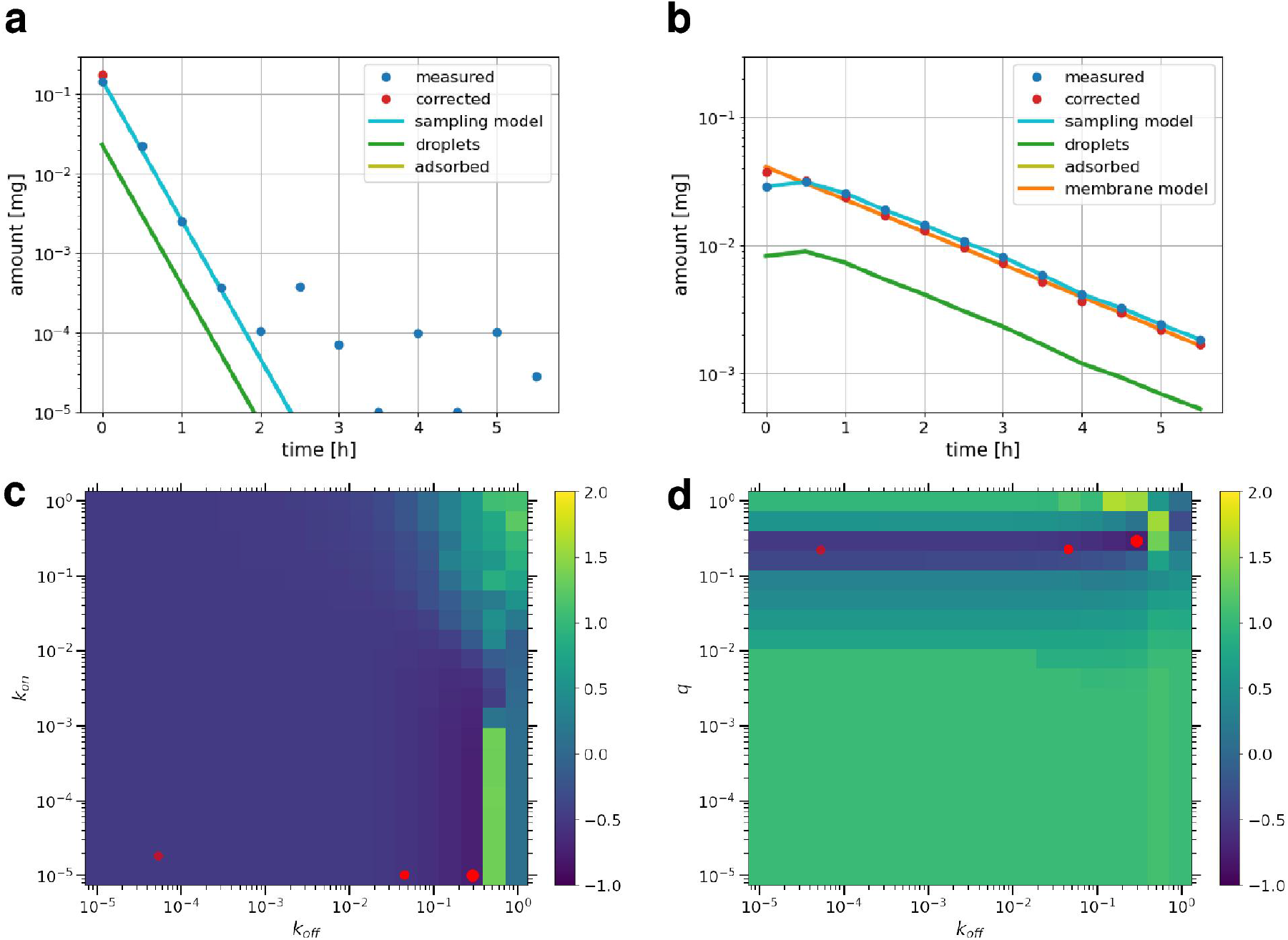
Sampling correction for FITC-dextran. Panels (a)-(d) are defined as in Fig. 3. The resolution limit of the optical spectroscopy used to quantify FITC-dextran corresponds to a concentration of approximately 10^−4^. Accordingly, values at or below this threshold in Fig. 4 should be interpreted as background.

For each analyte, the raw measured amounts *S* and the sampling-corrected inflow sequences *I* are shown under both experimental conditions: passive diffusion across an inert membrane (panels a) and sampling calibration in the absence of the transwell (panels b). Panels a−b of Fig. 3 (chloroquine) and Fig. 4 (FITC-dextran) display the measured sample sequences alongside the corrected inflow sequence (subsequently referred to as the corrected sample sequence), the best-fitting sampling model, and the predicted amounts retained in residual droplets (*x*) and adsorbed to surfaces (*y*). Both Figs. 3b and 4b include the fitted two-compartment diffusion model. Uncertainty in the sampling correction is visualized through the spread of individual traces corresponding to jackknife estimates of the sampling parameters [21]. Panels c– of both Fig. 3 and Fig. 4 present heatmaps of the fitting error Δ_total_ over two planar sections of the sampling parameter space that include the optimal parameter set. Overlaid points show the projections of the jackknife parameter estimates into these planes.

To reflect the practical detection limits of the analyte, we modify the error metric used during fitting (originally defined in Eq. (5)) by applying a lower cutoff, *x*_min_, to both observed and predicted values:

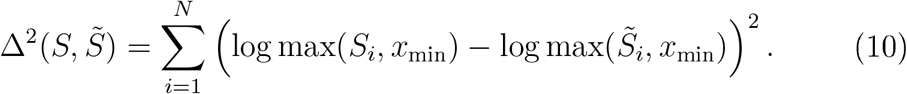

The sampling model predicts substantial surface adsorption for chloroquine and negligible adsorption for FITC-dextran. For both compounds, residual droplet retention accounts for approximately 20% of the analyte when the transwell insert is present – a contribution that includes fluid remaining underneath the insert during sample aspiration. The parameters for chloroquine are well constrained, as indicated by the sharp global minimum in the heatmaps and the tight clustering of the jackknife parameter estimates (Fig. 3c, d). In contrast, the adsorption parameters for FITC-dextran are less well defined: a broad range of sufficiently small *k*_On_ values yields fits of comparable quality, and for small *k*_On_, the fit becomes largely insensitive to *k*_Off_ (Fig. 4c, d). Despite this ambiguity, the correction results are robust, jackknife parameter estimates are visually indistinguishable in Fig. 4a and b.

The calibration experiment for chloroquine in the absence of a membrane (Fig. 3a) exhibits a biphasic decay: an initial rapid decline followed by a slower phase. This behavior strongly supports the presence of two distinct retention processes—adsorption to surfaces and retention in residual droplets—each governed by its specific kinetics. For chloroquine, the sampling model predicts that adsorption significantly distorts the apparent decay rate in the sample sequence Fig. 3b). Sampling correction notably increases the estimated inflow for both analytes in the first sampling interval (Figs. 3b, 4b). The close agreement between the corrected sample sequences and the two-compartment model predictions, as well as the ability of the sampling model to reproduce the biphasic decay seen in membrane-free calibration experiments, supports the validity of the proposed data correction pipeline in disentangling true transport dynamics from sampling artifacts. Building on this framework, we next examine analyte transport across cell-covered membranes, where biologically relevant processes such as cellular uptake and metabolism also shape the sample profile.

### 3.2. Passive transport through cells

#### 3.2.1. The three compartment model

To account for passive diffusion through a cellular barrier layer atop a porous membrane, we extend the two-compartment model by introducing a third, cellular compartment, where the free analyte concentration is denoted by *c*_*C*_. The total analyte content within this compartment, *M*_*C*_, is assumed to be proportional to *c*_*C*_:

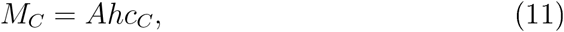

where *Ah* is the effective volume of the barrier cell layer. As suggested by others [10, 22], parameter *h* is a measure of reversible sequestration capacity within the cell layer: it reflects both the physical volume of the cells and the analyte-binding capacity of intracellular structures.

The time evolution of the system is governed by analyte fluxes *J*_*A*_ and *J*_*B*_, representing influx from the apical compartment (Eq. (6)) and efflux into the basolateral compartment (Eq. (7)), respectively. The mass balance for the cellular compartment is given by

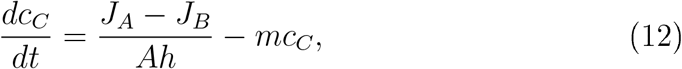

where *m* accounts for irreversible loss of analyte within the cell layer, such as metabolic degradation or covalent binding [22].

The apical flux into the cellular compartment is described by

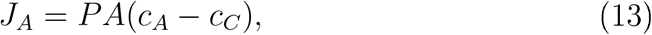

where *P* is the permeability of the apical cell membrane.

At the basal interface, the analyte must traverse both the basal cell membrane and the porous support. Assuming that the interfacial volume between these layers is negligible, we impose continuity of analyte flux:

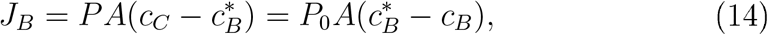

where *P*_0_ is the permeability of the porous support, and 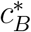

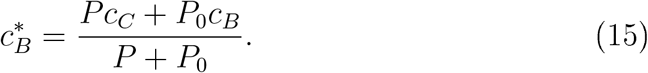

is the intermediate concentration at the basal interface. Solving Eq. (14) for 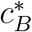 yields

Substituting this back into Eq. (14) gives the net efflux:

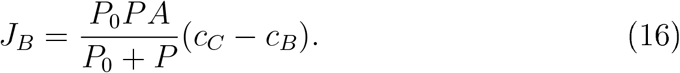

Together, Eqs. (6), (7), (12), (13), and (16) define the three-compartment model of passive transport across a cell-covered membrane. Several parameters (*A, V*_*A*_, *V*_*B*_, *P*_0_) are fixed by the experimental configuration; the remaining three, *θ*_*cell*_ = {*P, h, m*}, can be inferred by fitting the model to corrected inflow sequences. For this purpose, model simulations of sampling measurements are used as described in Section 3.1.3. At the beginning of each simulation, the apical, cellular, and basolateral concentrations are initiali ed to *c*_*A*_(0) = *c*_0_, *c*_*C*_(0) = 0, and *c*_*B*_(0) = 0, respectively. After each sampling event, the basolateral concentration is reset to zero, reflecting the medium replacement that occurs during the experimental sampling process.

Figure 5a shows simulated inflow sequences for a range of permeability values *P*, assuming no sequestration or degradation (*h* = 0, *m* = 0). In this limit, the model predicts simple exponential decay, with a time scale determined by *P*.

**Figure 5.**
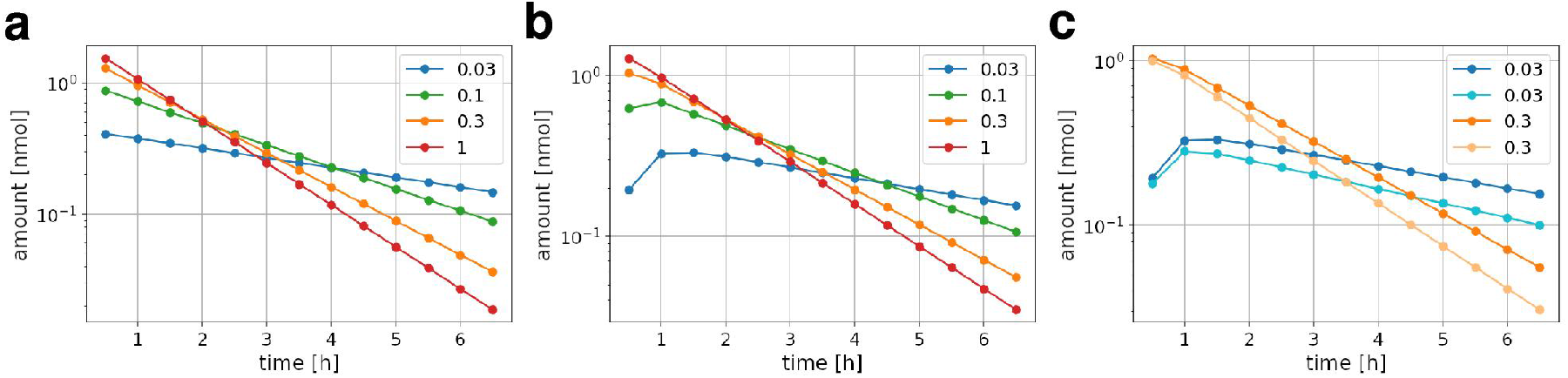
Effects of the three-compartment model parameters: permeability *P* (a), effective cell layer height *h* (b), and metabolic loss rate *m* (c). a: Predicted time course of analyte amounts in the receiver compartment for a negligible metabolic rate (*m* = 0) and a thin barrier layer (*h* = 0.01 mm) and four values of permeability: *P* = 0.03, 0.1, 0.3, and 1 mm/min, as indicated in the legend. Membrane permeability was set to *P*_0_ = 0.6 mm/min. b: The effect of barrier thickness on analyte transport. The effective cell layer height is increased to *h* = 1 mm, with the same set of *P* values as in panel (a). c: For two permeability conditions (*P* = 0.03 and 0.3 mm/min), analyte profiles are shown for *m* =0 (darker curves) and *m* = 0.1 (lighter curves), with *h* = 1. Metabolic loss reduces the accumulation rate of the analyte in the receiver, with stronger effects at lower permeability values.

Figure 5b illustrates the effect of reversible sequestration by setting *h* = 1 and varying *P*. Sequestration acts as a buffer, delaying the appearance of analyte in the basolateral compartment and suppressing early accumulation, particularly at lower *P*. This buffering can reduce the analyte level in the first sample and shift the peak to later time points.

Figure 5c demonstrates the effect of irreversible loss (*m>* 0) by comparing simulations with *m* = 0 and *m* = 1*/h*, using an intermediate value of *h*. As *m* increases, the decay of the sample sequence becomes more pronounced, reflecting progressive loss during transcellular passage.

#### 3.2.2. Parameter estimation from cellular barrier transport data

To analyze transport across cell layers, we employ a two-step procedure that builds upon the above-established sampling correction and model fitting methods (Fig. 6). In the first step, the measured sample sequences *S* are corrected using known sampling distortion parameters *θ*_*sampling*_, yielding corrected inflow sequences *I*. These sequences represent the amount of analyte entering the basolateral compartment between sampling points, with distortions due to droplet retention and surface adsorption removed.

**Figure 6.**
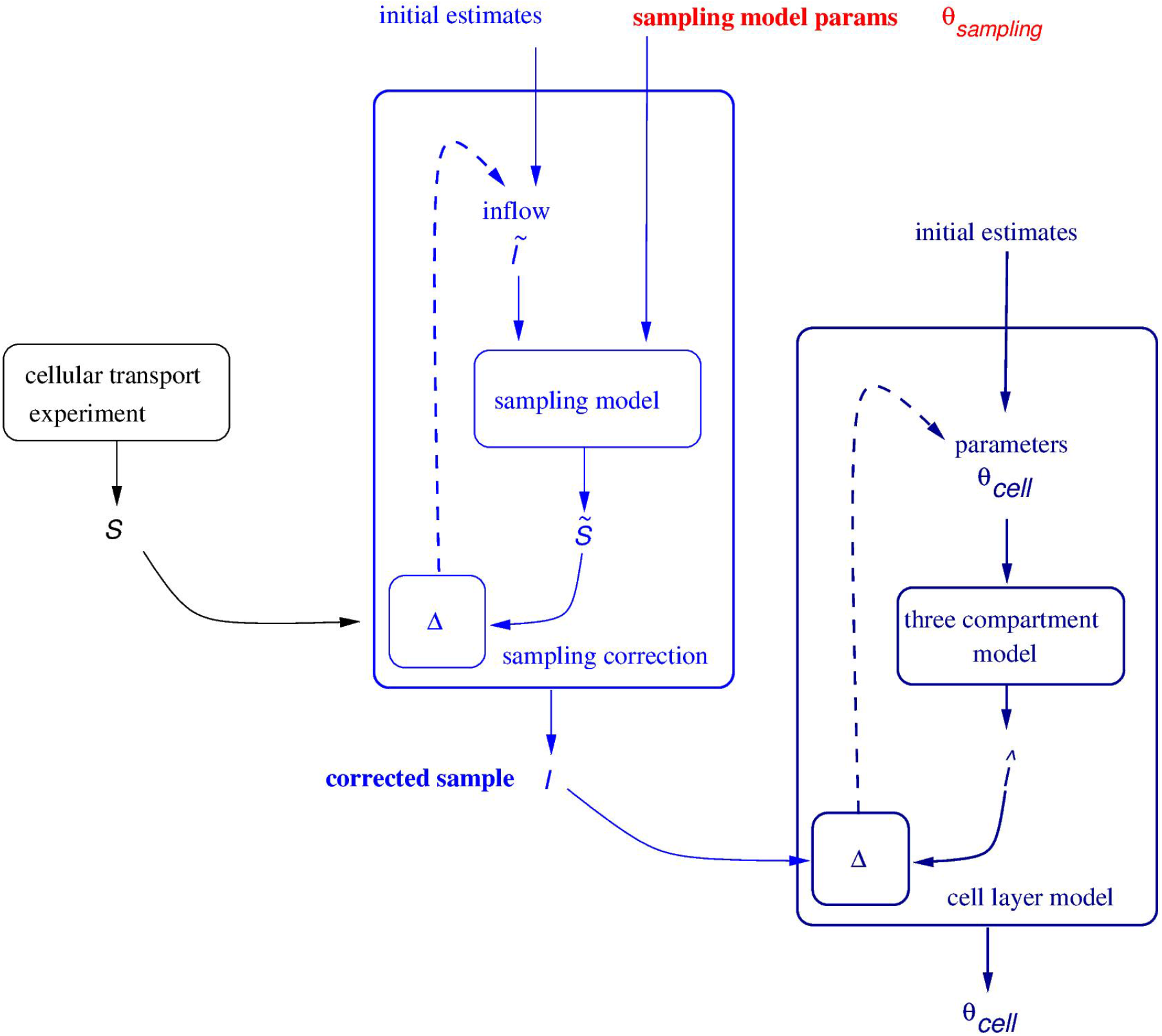
Two-step fitting procedure for estimating three-compartment model parameters from transwell barrier cell culture transport experiments. As in Fig. 2, algorithmic blocks with inputs and outputs are shown as boxes and arrows; dashed lines indicate optimization steps. First, the sampling model (blue) is used to correct measured sample sequences *S* (black), yielding the corrected inflow sequences *I*. These inflow sequences are then used to fit the three-compartment cellular transport model (dark blue). The discrepancy between *I* and the model-predicted inflow sequences *Î* is minimized to determine the best-fit parameter set *θ*_*cell*_.

In the second step, the corrected inflow sequences *I* are fitted using the three-compartment model of cellular transport. Simulated inflow sequences *Î* are generated based on trial values of the cellular transport parameters *θ*_*cell*_. The fit quality is quantified using the logarithmic deviation metric Δ^2^(*I, Î*). During fitting, the membrane permeability *P*_0_ is held fixed at the value determined from cell-free calibration experiments, while the parameters *θ*_*cell*_ are optimized to minimiz e Δ^2^.

As an example, Fig. 7 shows both the measured and corrected sample sequences for a data set taken from an experimental study [20]. The study involved chloroquine transport across a Caco-2 cell layer grown on a transwell insert. The figure also displays the predicted amounts of chloroquine retained in residual droplets, along with a substantial fraction adsorbed to surfaces. Notably, the correction reduces the apparent delay in the concentration peak, and the three-compartment model provides a good fit to the corrected data. While these results suggest that the fitting process can capture key features of cellular transport, we next turn to evaluating the reliability and limitations of parameter inference.

**Figure 7.**
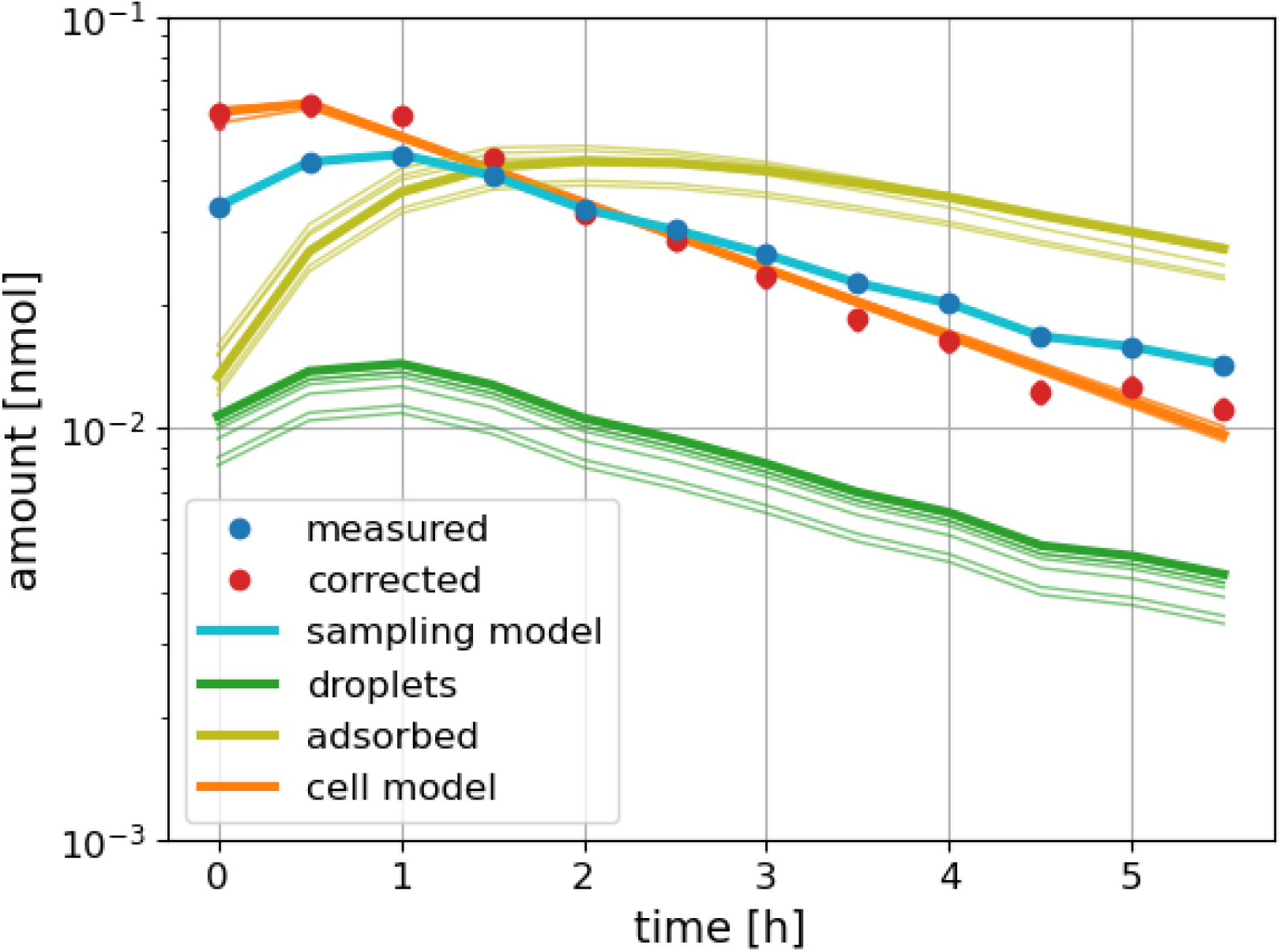
Chloroquine transport across a Caco-2 monolayer. Measured sample sequence (blue), corrected inflow sequence (orange), and fitted sampling model (green) for membrane-free calibration experiments. Also shown are the predicted amounts retained in residual droplets (red) and adsorbed to surfaces (magenta), with jackknife uncertainty indicated by shaded traces. The brown line shows the best-fit three-compartment model of passive transport across a cellular barrier layer.

The three-compartment model predicts analyte transport as a function of permeability (*P*), reversible sequestration (*h*), and metabolic loss (*m*). However, these parameters are not always uniquely identifiable from experimental data: In some regions of the parameter space, certain parameters become irrelevant and thus unidentifiable, regardless of the fitting method. Even when the model is sensitive to all three parameters, the range of possible output behaviors is limited, and distinct parameter combinations can yield similar infiow profiles. To assess parameter identifiability, we simulated the model over a 3D grid of parameter sets *θ*_*cell*_ = {*P, h, m*} spanning several orders of magnitude in each dimension. Pairwise dissimilarities between the resulting infiow sequences 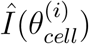 and 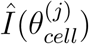 were quantified using the logarithmic deviation metric Δ^2^.

Figure 8a evaluates the sensitivity of the model to the parameter *P*. For each point in the heatmap, we calculated the minimum Δ^2^ dissimilarity to any other simulation with a different *P* value:

**Figure 8.**
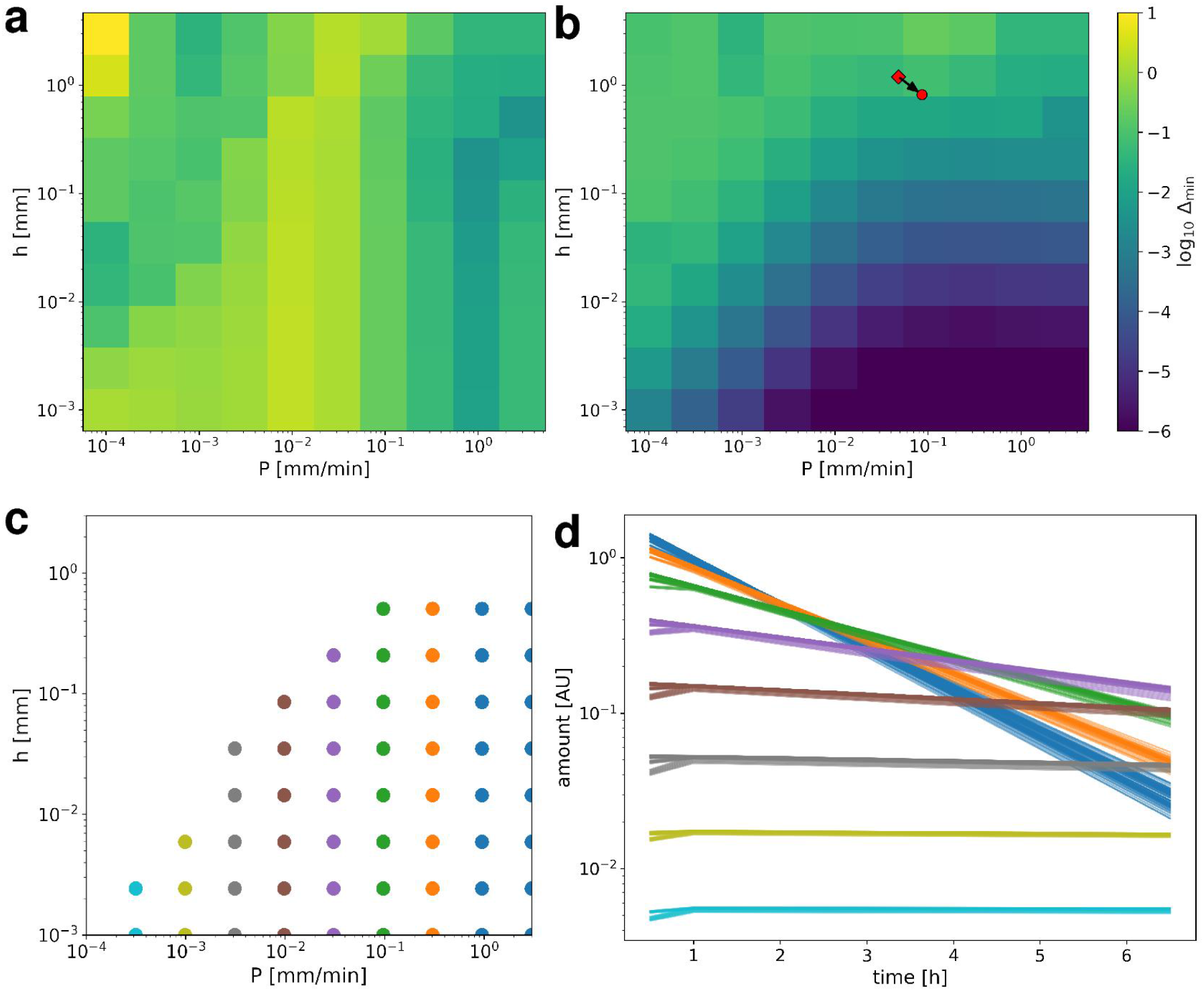
Identifiability analysis of the three-compartment model. (a) Sensitivity of model output to the permeability parameter *P*, measured by *δ*_*P*_: the minimum Δ^2^ distance between each simulated inflow sequence and any other with a different *P* value. Low values (dark) indicate poor resolvability of *P*. (b) Overall output diversity across the parameter space. The heatmap on the (*P, h*) plane shows the minimum Δ^2^ distance to any other simulation, evaluated across all tested values of *m*. Red markers indicate best-fit parameters for chloroquine transport across a Caco-2 monolayer (Fig. 7), based on corrected (diamond) and uncorrected (circle) data. (c) Clusters of parameter combinations producing equivalent inflow sequences, each colored distinctly. (d) Simulated inflow sequences for all members of each cluster in (c), colored accordingly. The similarity of traces within each group illustrates the equivalence of model outputs. and degradation becomes negligible due to short transit times. Under these conditions, a broad range of (*h, m*) values produce nearly indistinguishable inflow profiles, rendering these parameters unidentifiable.

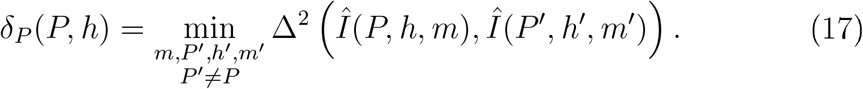

Low values of *δ*_*P*_ indicate regions where distinct *P* values yield similar outputs, requiring higher data precision for accurate inference. In particular, as *P* increases, the model output approaches the limiting behavior of a cell-free membrane. Additionally, high values of sequestration (*h*) can delay analyte inflow in a way that mimics reduced *P*, creating a second zone of limited resolvability.

Figure 8b expands the analysis to the full parameter set. Each point in the (*P, h*) heatmap represents the minimum dissimilarity between the corresponding simulation (across all tested values of *m*) and any other simulation:

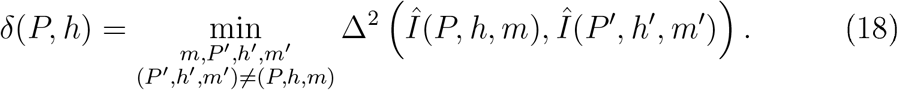

This global comparison captures output similarities arising from distant regions of parameter space. In general, resolving all three parameters requires substantially better precision than is needed to resolve *P* alone. In particular, when transcellular diffusion is rapid, sequestration has little effect,

To visualie such ambiguities, we defined a tolerance threshold *δ*_min_, set slightly above the lowest Δ-distance values observed in Figs. 8a-b. Simulations *i* and *j* were considered equivalent if Δ^2^(*Î*(*i*), *Î* (*j*)) *< δ*_min_. Figure 8c shows clusters in the *P* -*h* plane in which all simulations are equivalent, regardless of *m*. The corresponding inflow profiles are plotted in Fig. 8d. Outside these regions - specifically for intermediate *P* values and non-negligible sequestration or degradation - the model output is sensitive to all three parameters. In such cases, reliable parameter estimation is feasible, provided the sample sequence is sufficiently precise.

The importance of correcting for sampling artifacts is illustrated in Fig. 8b, where markers indicate best-fit parameter sets derived from both corrected and uncorrected sample sequences in a chloroquine transport experiment across a Caco-2 barrier layer (see Fig. 7). The fitted permeability *P* value increases by nearly a factor of two after correction. This shift reflects the fact that uncorrected sampling artifacts, such as delayed release and residual volume, can mimic intracellular retention. Without correction, the model compensates for the delayed signal by underestimating *P* and overestimating *h*. Accurate deconvolution of sampling distortions is therefore essential for reliable downstream parameter estimation.

## 4. Discussion

### 4.1 Relation to Apparent Permeability Measurements

The apparent permeability coefficient (*P*_app_) is widely used to quantify transcellular or transepithelial transport rates across biological barriers [23, 24, 13]. It is defined by Fick’s law, which describes solute tlux across an *inert* cell monolayer driven by a concentration gradient:

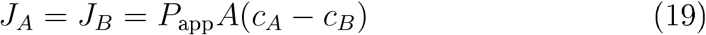

This expression is mathematically equivalent to the two-compartment model used in our membrane-only simulations and predicts an exponentially decaying sequence of analyte amounts in the sampled basolateral medium over time (see Figs. 3, 4). However, as shown in Fig. 7, early-phase sample concentrations can deviate substantially from a pure exponential decay. We attribute this to reversible interactions with the cell layer, including binding and sequestration, captured in our three-compartment model by the parameter *h*. After this transient phase, concentrations tend to follow an exponential decline, which often forms the empirical basis for *P*_app_ estimation [25]. Model simulations show, however, that the decay rate is not determined by cell layer permeability *P* alone, but also infiuenced by *h* and *m* (Fig. 5).

In this study, we suggest replacing *P*_app_ with the three parameters of a minimal three-compartment model because it reproduces observed transport profiles more accurately than Eq. (19), yet it remains simple enough for practical parameter estimation. The effective permeability *P* aggregates all transport routes across the cell layer, including transcellular, paracellular, and pericellular pathways [22, 26]. Importantly, *P* excludes the permeability of the porous support (*P*_0_) and effects such as unstirred medium layers [27], which are addressed separately via calibration using cell-free inserts. Figure 8 demonstrates that the proposed model parameters can be reliably estimated from data, provided they influence the observable outputs under the experimental conditions and measurement precision is adequate.

Extended models with additional mechanistic detail have been proposed in the literature [10, 22, 28, 29, 30]. However, the added complexity requires targeted experimental perturbations, such as transporter inhibition or fiow reversal, as the additional parameters are not identifiable from unperturbed sample sequences alone. When such targeted experimental data are available, these models can provide more detailed pharmacokinetic insight [31, 32]. However, when analysis is limited to sample sequences from routine, unidirectional transport experiments, we propose that the parameter triplet *θ*_*cell*_ = {*P, h, m*} captures the dominant dynamics more effectively than the traditional *P*_app_ value.

In many studies, *P*_app_ is calculated from the cumulative amount transported at a single time point *T*, yielding a time-specific coefficient 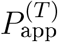. To assess how well this metric refiects underlying transport dynamics, we simulated barrier transport using the three-compartment model and computed 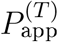 at *T* = 3 hours. Figure 9a shows a heatmap of 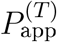 across a grid of *P* and *h* values with *m* = 0. As expected, 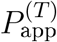 increases with *P*, but is suppressed by increasing *h*, which delays analyte appearance in the basolateral compartment. Figure 9b shows the analogous trend across the *h*−*m* plane at fixed *P* : both reversible sequestration and irreversible loss reduce transport even when *P* is unchanged.

**Figure 9.**
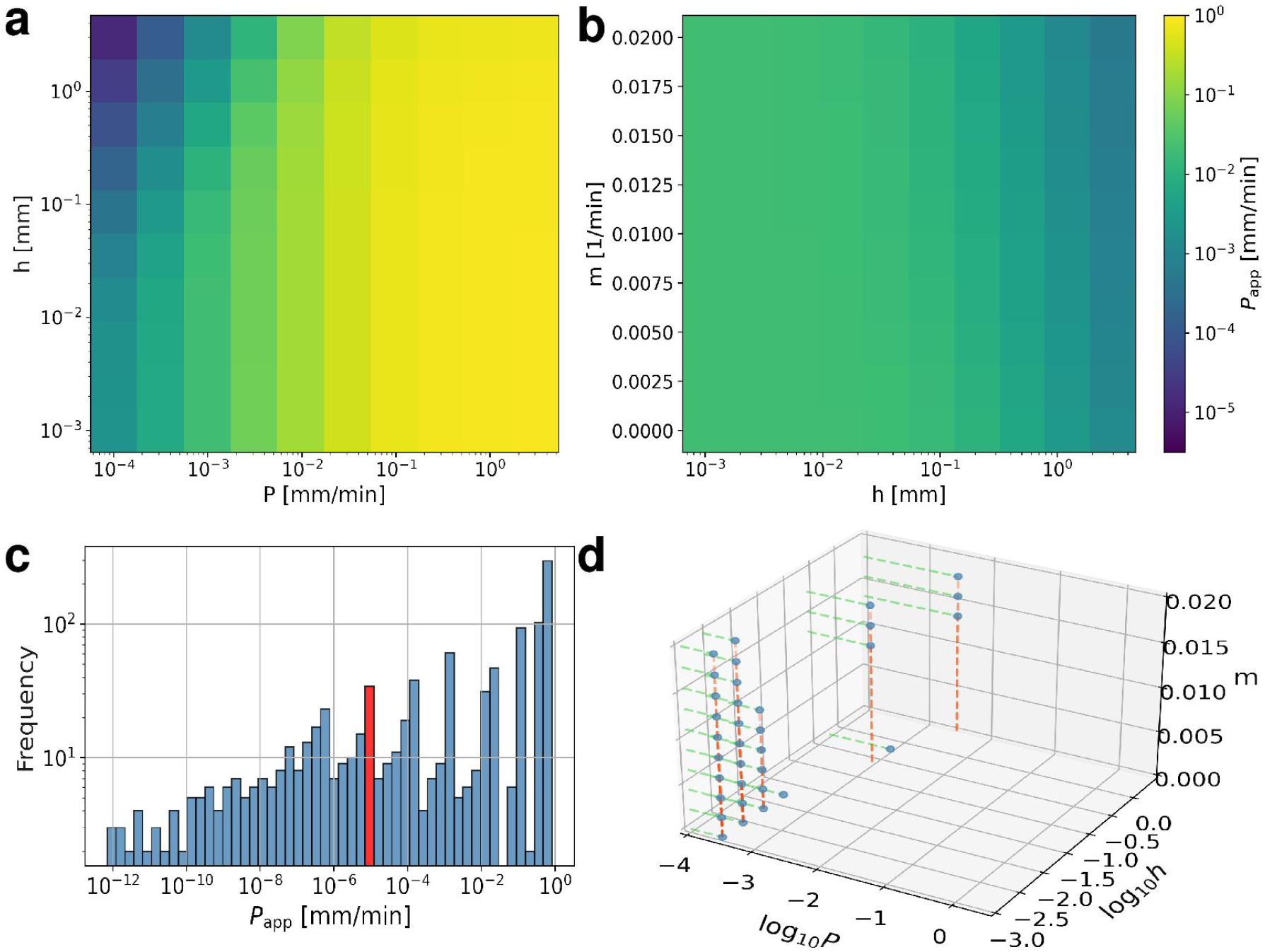
Influence of three-compartment model parameters on the apparent per-meability coefficient 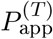, evaluated at *T* = 3 hours. a: Heatmap of 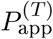 values as a function of membrane permeability *P* and effective cell layer height *h*, under conditions of no metabolic loss (*m* = 0). b: Similar heatmap for fixed *P* = 10^*−*3^ mm/min, across a range of *m* and *h* values. The membrane permeability is held constant at *P*_0_ = 0.6 mm/min. c: Histogram of 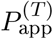 values obtained from simulations spanning a wide range of (*P, h, m*) parameter combinations. d: Subset of (*P, h, m*) parameter combinations that lead to the specific 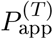 value high-lighted in red in panel (c). These parameter sets form a manifold in the model’s three-dimensional parameter space, underscoring the challenges of inferring unique transport properties from a single apparent permeability measurement.

As a different perspective, the histogram in Fig. 9c displays the spread of 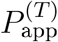 values from all simulations. A narrow target range of 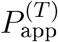 values (marked in red) corresponds to a broad set of underlying parameters, as illustrated in the scatter plot in Fig. 9d. These findings confirm that while *P* is the primary determinant of *P*_app_, the apparent permeability is modulated, often by an order of magnitude, by both *h* and *m*. These results underscore the limitations of relying on single-point estimates. Full time-course measurements, combined with mechanistic models of balanced complexity, offer a more accurate and interpretable characteriation of analyte−cell interactions.

### 4.2. Improvements in Sampling Correction and Transport Modeling

Building on our earlier work [16], where we introduced a mechanistic framework to correct sampling artifacts in automated membrane transport systems, we now present a refined methodology that improves both accuracy and applicability. The recursive linear model for sampling distortion (Eqs. 1–4) retains its original structure but is applied under improved experimental conditions: placing the culture vessel on a rotary shaker ensures thorough mixing, allowing us to treat the basolateral compartment as well-mixed and eliminate the need for internal sub-compartment modeling.

A key methodological advance is the introduction of a nested two-stage fitting procedure. Sampling parameters are first inferred from membrane-only experiments, and then used to correct sample sequences from cellular barrier measurements. Mechanistic transport models are subsequently applied to the deconvolved data. This separation avoids conflating technical artifacts with biological transport phenomena and enables more reliable parameter estimation. The importance of these corrections is illustrated by our analysis of both FITC-dextran and chloroquine. In the former, a hydrophilic macromolecule with minimal surface binding, distortion arose mainly from droplet retention. In contrast, chloroquine revealed strong surface adsorption effects, with substantial analyte accumulation on tubing or vial walls. These artifacts, if uncorrected, would lead to underestimation of permeability (Fig. 8b) and misinterpretation of the underlying mechanisms.

The recursive correction model accounts for both residual volume and reversible adsorption. Its robustness is reflected in the reproducibility of the fitted sampling parameters across experiments. Because many automated sampling platforms share similar physical characteristics–tubing networks, dead volumes, and sequential collection–the same framework can likely be adapted to other systems with minimal changes.

To model analyte transport across a cell-covered membrane, we used a minimal three-compartment model incorporating three interpretable parameters: *P* (effective permeability of the cell layer), *h* (reversible sequestration capacity), and *m* (irreversible loss rate). The parameter *h* captures in a single linear approximation the rapid, reversible interactions between the analyte and cellular components, such as intracellular binding, partitioning, or temporary exclusion. It is conceptually related to earlier partition models [10, 33]. The term *m* accounts for metabolic degradation or irreversible binding during transcellular passage [22]. While all components of the model have been proposed before, we emphasi’.e its use here as a minimal but sufficient model that can reproduce the sample sequences observed in multiple transport experiments.

This capability has direct implications for pharmaceutical and preclinical research. Transwell assays are widely used to predict absorption, intestinal permeability, and blood-brain barrier penetration. Misestimation of passive permeability due to uncorrected sampling artifacts or oversimplified models can affect downstream pharmacokinetic predictions. By fitting a mechanistically grounded model to deconvolved data, our framework provides more accurate estimates of the cellular transport parameters most relevant to drug disposition.

### 4.4. Relevance of transwell-based cellular barrier assays

Transwell-based assays are a cornerstone of in vitro permeability testing, offering a practical compromise between experimental control and biological relevance. When applied to cell-covered membranes, these assays enable the investigation of compound interactions with cellular components, including transporters, receptors, and metabolic enzymes–factors critical to understanding drug disposition and potential drug–drug interactions [13]. Speciali’.ed cell lines and culture conditions further allow modeling of tissue-specific barriers, such as the blood–brain barrier or pulmonary epithelium.

Despite these advantages, cell-based permeability assays are associated with several challenges. Compared to artificial membrane systems 34, 35J, they involve higher costs due to the need for sterile facilities, skilled personnel, and ongoing culture maintenance. Throughput is also limited by the manual steps involved, and biological variability - arising from cell passage number, differentiation status, and minor protocol differences - can impair reproducibility [13]. Automated workflows, including the sampling system investigated in this study, can mitigate some of these limitations by reducing manual variability and enabling more consistent time-course data acquisition. The field continues to evolve toward more physiologically relevant in vitro systems. Emerging technologies such as 3D cell cultures, co-culture barriers, and organ-on-a-chip platforms aim to better replicate in vivo conditions by incorporating extracellular matrix components, dynamic f:ow, and multiple interacting cell types [36]. While these models offer enhanced realism, they also increase system complexity and further underscore the importance of robust analytical and modeling tools to interpret transport measurements across diverse experimental platforms. In this context, the core principles demonstrated in this study – sampling correction, model-based deconvolution, and minimal mechanistic parameterization – remain broadly applicable and may help bridge the gap between traditional assays and next-generation barrier models.

## Funding

This work was supported by the National Research, Development and Innovation Office, Hungary (grants OTKA ANN 132225 (A.C.) NKFIH K-135712 (I.B.), K-142904 (S.B.), VEKOP-2.3.3-15-2017-00020 (S.B.). BD was supported by the Austrian Science Fund (grant FWF I4677-B). Additional support was provided by the institutional grant EKA 2023/071-P085-1.

The funders had no role in study design, data collection and analysis, decision to publish, or preparation of the manuscript.

## Notes

https://github.com/aczirok/barrier-transport-tools

### Competing Interest Statement

A.C. and T-Z.J. are founders of BioPhys-Concepts KFT, a university start-up company that commercializes millifluidic technology.

